# ProMetheusDB: an in-depth analysis of the high-quality human methyl-proteome

**DOI:** 10.1101/2021.09.20.461082

**Authors:** Enrico Massignani, Roberto Giambruno, Marianna Maniaci, Luciano Nicosia, Avinash Yadav, Alessandro Cuomo, Francesco Raimondi, Tiziana Bonaldi

## Abstract

Protein Arginine (R) methylation is a post-translational modification involved in various biological processes, such as RNA splicing, DNA repair, immune response, signal transduction, and tumour development. Although several advancements were made in the study of this modification by mass spectrometry, researchers still face the problem of a high false discovery rate. We present a dataset of high-quality methylations obtained from several different heavy methyl SILAC (hmSILAC) experiments analysed with a machine learning-based tool doublets and show that this model allows for improved high-confidence identification of real methyl-peptides. Overall, our results are consistent with the notion that protein R methylation modulates protein:RNA interactions and suggest a role in rewiring protein:protein interactions, for which we provide experimental evidence for a representative case (i.e. NONO:PSPC1). Upon intersecting our R-methyl-sites dataset with a phosphosites dataset, we observed that R methylation correlates differently with S/T-Y phosphorylation in response to various stimuli. Finally, we explored the application of hmSILAC to identify unconventional methylated residues and successfully identified novel histone methylation marks on Serine 28 and Threonine 32 of H3.

## INTRODUCTION

Protein methylation is a widespread PTM that consists of the addition of one or more methyl (CH_3_) groups to a residue, most frequently an Arginine (R) or Lysine (K). Arginines can be mono-methylated (MMA) or di-methylated, and the di-methylation can be symmetrical (SDMA) if the two methyl groups are bound to different nitrogen atoms of the guanidino group or asymmetrical (ADMA), if they are bound to the same atom. Overall, methylation does not change the charge state of R, but alters their steric hindrance and reduces their ability to form hydrogen bonds with other molecules, thus affecting their interactions with other biomolecules, both positively and negatively (Fulton et al, 2019). Along this line, the distinction between ADMA and SDMA is crucial, because the two modifications present different steric hindrances, hence they are recognized and bound by distinct protein domains (readers) and can lead to completely different functional outcomes, as exemplified by asymmetrical and symmetrical di-methylation on R3 of histone H4 (H4R3) that are associated to transcriptional activation and repression, respectively (Wysocka et al, 2006).

The biological donor of methyl groups in the enzymatic reaction of protein methylation is S-Adenosyl-Methionine (SAM), which is synthesized by the enzyme methionine adenosyltransferase (MAT, most commonly known as SAM synthase) from Methionine (M) and ATP (Murray et al, 2014). Specifically, the transfer of methyl-groups to R is carried out by the Protein Arginine Methyltransferases (PRMTs) a class of enzymes consisting of nine proteins, classified in three groups: type I PRMTs (including PRMT1, PRMT2, PRMT3, PRMT4/CARM1, PRMT6, and PRMT8) catalyse the formation of MMA and ADMA; type II PRMTs (PRMT5 and PRMT9) catalyse the formation of MMA and SDMA; PRMT7, which is the only type III PRMT, leads to MMA only (Lorton & Shechter, 2019). PRMT1 is the most active enzyme among type I PRMTs and PRMTs in general, being overall responsible for ∼85% of the methylation events in the human cell (Yamaguchi & Kitajo, 2012), whereas PRMT5 is the most active type II PRMT (Stopa et al, 2015).

While methylation of histones and its implication in gene expression regulation has been extensively studied in several model systems and functional states (Di Lorenzo & Bedford, 2011), the concept that K/R methylation is widespread on various non-histone proteins involved in a variety of important biological processes is more recent and has been supported by several studies that have been accumulating in the last 10 years (Bremang et al, 2013; Geoghegan et al, 2015; Guccione & Richard, 2019; Larsen et al, 2016; Wu et al, 2021). In particular, PRMT1 is involved in DNA-Damage Response (DDR) by directing its activity towards chromatin proteins (such as RBMX, CHTOP and DDX17 (Musiani et al, 2020)) and DDR proteins such as MRE11 and 53BP1 (Vadnais et al, 2018). PRMT4/CARM1 regulates nonsense-mediated decay and pre-mRNA splicing (Sanchez et al, 2016), and PRMT7 is involved in stress responses such as heat and proteasome inhibition (Szewczyk et al, 2020). In addition, PRMT1, PRMT4, PRMT5 and PRMT7 have all been reported to catalyse methylation on RNA-binding proteins (RBPs), affecting their interactions with RNA molecules, with implications in the regulation of mRNA splicing, miRNA maturation, translation and RNP granules assembly (Cheng et al, 2007; Fong et al, 2019; Guccione & Richard, 2019; Maniaci et al, 2021; Schisa & Elaswad, 2021; Spadotto et al, 2020; Szewczyk et al, 2020).

Importantly, PRMTs are often aberrantly expressed in cancer, including both solid tumours (i.e., brain, breast, lung, colon, bladder, head and neck cancer) and haematological malignancies (such as leukaemia) (Bao et al, 2013; Hwang et al, 2021) and in other diseases like neurodegeneration (Couto et al, 2020) and metabolic disorders (vanLieshout & Ljubicic, 2019). Hence, there is a great interest in developing therapies that are based on the pharmacological modulation of these enzymes. In fact, several PRMTs inhibitors are under development (Hwang et al, 2021; Smith et al, 2018), with some already undergoing clinical trials (trial identifiers NCT03573310, NCT02783300 and NCT03614728).

Recent studies on protein methylation took advantage of the progress made by Liquid chromatography-tandem Mass Spectrometry (LC-MS/MS)-based proteomics technology. Several MS-based studies have led to the annotation of progressively larger methyl-proteomes; however, it has also been shown that the proteome-wide identification of methylation sites through LC-MS/MS is prone to high false discovery rates, due to the presence of artefacts isobaric to this PTM (Hart-Smith et al, 2016). For instance, the chemical methyl-esterification of an acidic residue (such as Aspartate (D) and Glutamate (E)) can be erroneously identified as an in vivo methylation on a nearby R residue by peptide search engines. Also, methylations are isobaric to some conservative amino acid substitutions, such as Alanine (A) to Valine (V), or D to E. To address these issues, the group of M. Mann proposed the heavy methyl SILAC (hmSILAC) metabolic labelling strategy (Ong & Mann, 2006), whereby cells are grown in presence of either natural Methionine (M0) or stable isotope-labelled [^13^CD_3_]-Methionine (M4) to isotopically label the methyl groups. Upon mixing the heavy and light samples, methylated peptides can be detected by MS as pairs of MS1 peaks separated by a specific mass difference; instead, chemical artefacts appear as single light peaks, since the heavy methyl groups can only be derived from an enzymatic reaction.

Until recently, hmSILAC use had been held back by the lack of computational solutions for the data analysis. Common peptide search engines, such as Andromeda or Mascot, can handle isotopic labelling of amino acids (as in traditional SILAC) but cannot be used when the label is encoded by a variable modification. To address this problem, our and M. Wilkins’s group developed two computation tools, tailored to process hmSILAC MS data for the identification of peptides methylated in vivo named hmSEEKER and MethylQuant, respectively (Massignani et al, 2019; Tay et al, 2018). Both tools have been successfully employed to expand the high-confidence annotation of the methyl-proteome in *Homo Sapiens* and *S. cerevisiae*, while tightly controlling the number of false-positive identifications (Geoghegan et al, 2015; Hamey et al, 2021; Musiani et al, 2019; Musiani et al, 2020).

Since the first release of hmSEEKER, we have improved our method, by adjusting the protocol for searching methyl-sites in MaxQuant and by upgrading hmSEEKER itself. Following a strategy similar to that described in (Massignani et al, 2019), we trained a Machine Learning (ML) model to recognize hmSILAC doublets generated by M-containing peptides, then we used the model to identify heavy-light methyl-peptides pairs. Thus, we were able to re-analyse all the hmSILAC experiments collected in our laboratory so far.

The majority of MS-based methyl-proteomics studies, including this one, have traditionally focused mainly on R and, to a lower extent, on K protein methylation, due to multiple reasons: first, from a technical point of view, there is a large variety of biochemical methods for the enrichment of peptides or proteins bearing these PTMs; second, the enzymes responsible for deposing or removing these PTMs are known, hence it is easier to correlate K/R methylation dynamics with the modulation of the respective enzyme, or enzyme family; third, a panel of small molecule inhibitors for the modulation of these enzymes is available. Despite this bias, there is little but increasing data showing that methylation naturally occurs also on other amino acids, such as D, E, Asparagine (N), Glutamine (Q), Serine (S), Threonine (T) and Histidine (H) (Afjehi-Sadat & Garcia, 2013). For instance, methylation of Q104 on histone H2A is enriched in the nucleolus compartment, where it is recognized specifically by RNA polymerase I (Tessarz et al, 2014). Concerning non-histone proteins, H methylation has been described on actin, myosin, MLCK2 and RPL3 (Kwiatkowski & Drozak, 2020), with suggested its involvement in muscle contraction.

Overall, our proteomics and computational analysis produced what, at the time of writing, is the largest orthogonally validated methyl-proteomics dataset. This dataset has been subsequently integrated with a list of R-sites that are significantly regulated in perturbed SILAC experiments, generating the ProMetheus database (ProMetheusDB) of high-confidence protein methylation sites.

We then have carried out an extensive functional analysis of the ProMetheusDB, confirming the role of R methylation in different aspects of RNA metabolism, but also suggesting its involvement in novel cellular processes, such as antigen processing and presentation, macrophage metabolism and immune response. We also investigated the potential cross-talk between R methylation and S/T-Y phosphorylation and carried out a structural analysis of methylated Rs at 3D interaction interfaces, which pointed towards a role of R methylation in fine-tuning protein:protein interactions. In this context, we focused on NONO protein, a target of PRMT1, and its interactor, the paraspeckle component PSPC1, showing that-indeed-R methylation of NONO directly impacts on its binding with PSPC1.

## RESULTS

### Implementation of machine learning in hmSEEKER v2.0 to improve the detection of methyl-peptides doublets from hmSILAC data

During the first implementation of hmSEEKER (Massignani et al, 2019), we defined empirical cut-off values to distinguish true hmSILAC doublets from false positives. However, these criteria were conservative and resulted in a very low False Discovery Rate (FDR) but also a suboptimal sensitivity. We addressed this point by training a machine learning logistic regression model to discriminate between putative true and putative false hmSILAC doublets (Fig 1A). This model allowed us to increase the number of hmSILAC-validated methyl-peptides in our dataset without compromising the FDR and provides more rigorous criteria for the identification of hmSILAC doublets compared to the original ones, which were estimated empirically. To obtain a dataset for model training, we followed the same strategy employed in (Massignani et al, 2019) to determine optimal cut-offs. Specifically, we analysed MS raw files from hmSILAC experiments to identify non-methylated peptides that contained one or more Methionine (M) residues: in fact, the labelling with M0 (“light”) or M4 (“heavy”) causes M-containing peptides to generate hmSILAC doublets that have the same properties as those generated by methyl-peptides. Thus, we used M-containing peptides as “mock methyl-peptides”. To model True Negatives, we also included in the dataset peptides without M and randomly assigned methylations to them, to mimic an erroneous identification by MaxQuant (Fig 1B). The peptides in the dataset were processed with hmSEEKER using the following cut-offs: |Mass Error (ME)| < 100 ppm, |Retention time difference (dRT)| < 25 and |log_2_ H/L intensity ratio (LogRatio)| < 25. These cut-offs allowed us to retrieve as many putative doublets as possible, to have enough data for the training of the model. Doublets generated by a “mock methyl-peptide” were then filtered to only include “Matched” doublets (i.e. those where both the heavy and the light peptides are identified, which are the most confident) and labelled “1” (n = 4434); doublets generated by a “true negative” peptide were labelled “0” (n = 3618). In total, the dataset used to train the predictor consisted of 8052 doublets.

**Figure 1.**
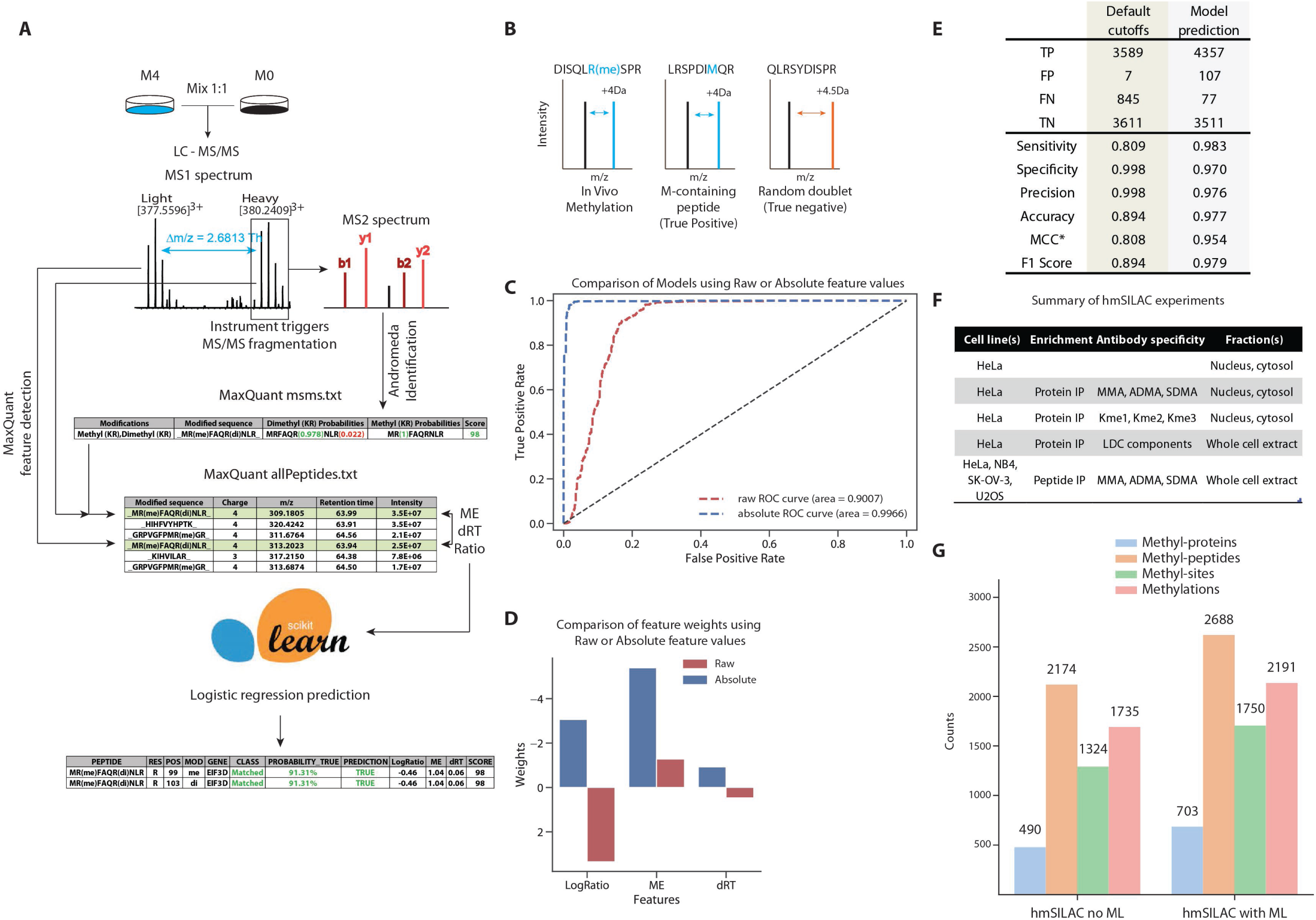
Rationale of hmSEEKER, development of the Machine Learning model and analysis of hmSILAC data. **A)** Schematic representation of the hmSEEKER workflow upon metabolic labelling with stable-isotope encoded Methionine (M). Cells grown in “light” (M0) and “heavy” (M4) media up to 10 passages are mixed in 1:1 proportion, then proteins are extracted, digested with proteases and analysed by LC-MS/MS. MaxQuant detects MS1 peptide peaks in MS raw data. Peaks with an associated MS2 fragmentation spectrum are processed by the database search engine Andromeda, to obtain peptide and PTM identifications. The hmSEEKER software reads MaxQuant peptide identifications and, for each methyl-peptide that passes the quality filters, finds its corresponding MS1 peak, then searches the corresponding heavy/light counterpart. A peak doublet is defined by the difference in the retention time (dRT), the log-transformed intensity ratio (LogRatio) and the deviation between expected and observed m/z delta (Mass Error, ME); these 3 parameters are used to predict if the peak pair is a true hmSILAC doublet or a false positive. **B)** Schematic examples of true positive and true negative doublets. Our rationale for training the machine learning model was that M-containing peptides produce hmSILAC doublets that are indistinguishable from those generated by methyl-peptides. **C)** Receiving Operator Characteristics (ROC) curves obtained by testing the models trained by using either the raw or absolute value of the features. **D)** Representation of the ML model weights with (blue) and without (red) taking the absolute values of the features. **E)** Comparison of the performance of hmSEEKER with (New ML Model) and without (Old Cut-offs) the ML predictor (*MCC = Matthews correlation coefficient). **F)** Summary of hmSILAC experiments analysed to produce the orthogonally validated methyl-proteome. A detailed description of these experiments is available in Table EV1. **G)** Composition of the high-confidence hmSILAC R-methyl-proteome before (left) and after (right) the implementation of the ML model.

The independent variables (features) used within the logistic regression model were the ME, dRT and LogRatio; the dependent variable was whether the doublet was a true hmSILAC doublet (“1”) or a random peak pair (“0”). The model was trained on 6000 doublets (3000 true + 3000 false; Appendix Fig S1A) using stratified five-fold cross-validation. During the training process, we also tested transformations of the features. For instance, to reduce the impact of potential outliers, we applied a quantile transformation to the features, which were then normalized based on their median and IQR values. Moreover, we observed that taking the absolute values of the features improved the performance of the model, allowing it to reach an area under the ROC curve >0.99 (Fig 1C). Upon validation of the model on the remaining 2052 doublets, we found that less than 2% of the doublets were incorrectly labelled by the logistic regression (Appendix Fig S1B). Inspection of the logistic regression coefficients revealed that ME was the most important feature in distinguishing true and false doublets as it had the largest absolute weight (−5.393), followed by LogRatio (weight = −3.067), whereas dRT (weight = −0.927) appeared to be the least important (Fig 1D).

We then tested whether the inclusion of the ML model improved hmSEEKER v1.0 performance by comparing it to the default cut-offs defined in (Massignani et al, 2019): the entire set of 8052 doublets used for the training of the model was reanalysed twice, first with the default cut-offs (i.e. |ME| < 2 ppm, |dRT| < 0.5 min, |LogRatio| < 1) and then by using the model predictions. The logistic regression showed an increase in sensitivity and accuracy, with only a very limited reduction in specificity and precision (Fig 1E); the metrics F1 score and Matthew’s Correlation Coefficient (MCC) were also improved upon applying this model. Overall, these results prove our initial feeling that the initial cut-offs were very conservative.

### Re-annotation of the largest high-quality human methyl-proteome so far

We first applied hmSEEKER v1.0 as described in (Massignani et al, 2019) to our hmSILAC data. Briefly, the data consisted of samples from four different cell lines (HeLa, SK-OV-3, NB4 and U2OS) which were subjected to different biochemical pipelines prior to LC-MS/MS, such as immuno-enrichment of methyl-peptides with CST PTMScan kits or immuno-enrichment of proteins with antibodies against methyl-R, methyl-K or LDC components. These enrichment methods were combined with separation techniques such as polyacrylamide gel electrophoresis, isoelectric focusing and HpH-RP liquid chromatography to reduce sample complexity and further boost the identification of methyl-peptides (Fig 1F and Table 1). Samples were then acquired on a Q Exactive orbitrap instrument and the resulting MS raw data were processed with MaxQuant to identify R- and K-methyl-sites (as described in the Material and Methods section).

The high-confidence methyl-proteome obtained from hmSEEKER v1.0 contained 2174 methyl-peptides, mapping to 490 different proteins, which carried a total of 1324 methyl-sites and 1735 methylation events (Fig 1G, left). By re-analysing the hmSILAC dataset with hmSEEKER v2.0, we annotated 2688 methyl-peptides mapping on 703 proteins and 2191 methylation events distributed on 1750 methyl-sites (Fig 1G, right). Overall, we were able to identify 23% more methyl-peptides, 32% more methyl-sites and 26% more methylation events. This represents the largest, orthogonally-validated methyl-proteome annotated so far from human cancer cell lines.

Upon the analysis of the methyl-proteome dynamics upon different perturbations through the combination of the biochemical workflow for methyl-peptide separation and enrichment with standard SILAC metabolic labelling, we set to expand further this experimental dataset by including the list of methyl-peptides that resulted as significantly up- or down-regulated in at least one of the SILAC methyl-proteomic experiments acquired, whereby dynamic changes of methyl-site were profiled quantitatively upon different perturbations (e.g. a drug treatment or the modulation of a PRMT enzyme) (Fig 2A). The inclusion of these methyl-peptides in the hmSILAC repository was performed under the assumption that methyl-peptides displaying significant changes upon a specific biological stimulus are likely to be enzymatic, whereas amino acid substitutions and chemical artefacts introduced by sample preparation should remain unaffected by any type of perturbation. Specifically, the SILAC experiments from which the significantly regulated methyl-peptides were extrapolated from a set of experiments carried out by our group in previous studies, where R methylation changes were profiled in response to: 1) PRMT1 over-expression or knock-down (Spadotto et al, 2020); 2) treatment of cells with the PRMT5 inhibitor GSK591, or with the PRMTs type I inhibitor MS023 (Fong et al, 2019; Musiani et al, 2019); 3) treatment of cells with cisplatin (CDDP), a chemotherapy drug that induces PRMT1 accumulation in chromatin, thus inducing reduced R-methylation level of cytoplasmic proteins (Musiani et al, 2020). We thus obtained a final dataset consisting of 2865 methyl-peptides, 723 methyl-proteins, 1814 methyl-sites and 2271 methylation events (Fig 2B, left), which we named ProMetheusDB, a repository of high-quality MS-based identified K/R-methyl-peptides from human cancer cell lines.

**Figure 2.**
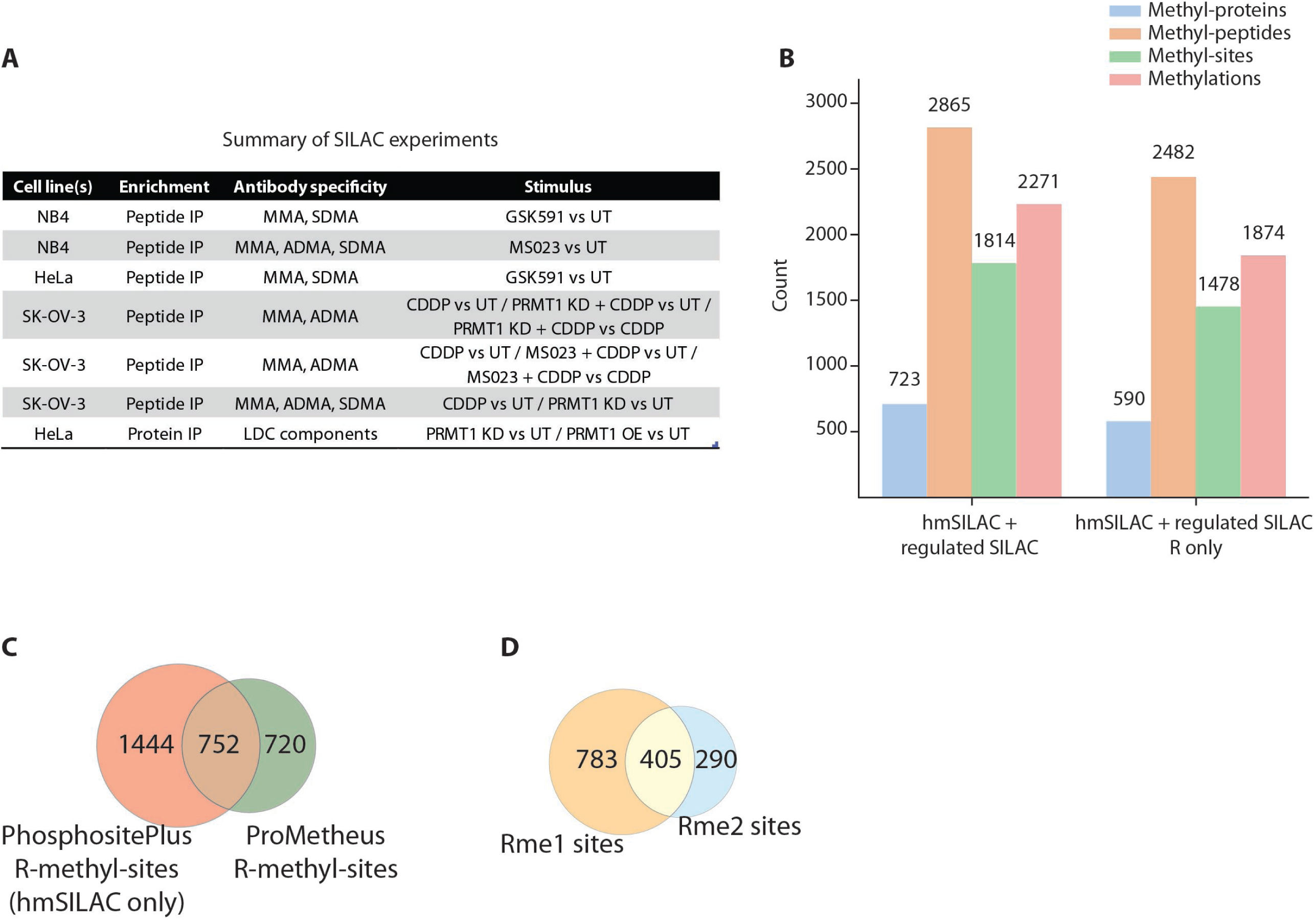
Integration of the hmSILAC and SILAC data. **A)** Summary of SILAC experiments used to expand the high-confidence methyl-proteome. Only peptides that were significantly regulated at least one experiment upon any of the perturbations used were included, based on the assumption that dynamically regulated peptides are unlikely to be artefact. A detailed description of these datasets and the linked experiments is available in Table EV1. **B)** Composition of the R-methyl-proteome upon the inclusion of the regulated SILAC methyl-peptides, representing the ProMetheusDB. **C)** Venn diagram comparing R-methyl-peptides in ProMetheusDB and annotated in Phosphosite Plus. **D)** Venn diagram comparing R-methyl-sites identified as mono-or di-methylated.

To characterise its features, we first checked its composition in terms of K and R methylation events. We identified proteins bearing K-methyl-sites or combinations of K- and R-methyl-sites (Appendix Fig S2A), although peptides on which K and R methylation coexist were overall quite rare (only 88, Appendix Fig S2B), thus suggesting that K- and R-methyl-sites are not necessarily neighbouring, but rather in different protein regions. It is worth noting that, while K methylation was included in the MS data analysis to avoid erroneous assignment of R methylation events to K and vice versa, we expected the coverage of the R-methyl-peptides in ProMetheusDB to be much larger than that of K-methyl-peptides, due to the employment of anti-pan-methyl-R antibodies in most of our experiments. Hence, we focused on the R-methyl-proteome, which comprises 2482 R-methyl-peptides, 590 proteins, 1472 R-sites and 1874 R-modifications (Figure 2B, right).

We first compared the R-methyl-sites in ProMetheusDB to the hmSILAC-validated R-methyl-sites in PhosphositePlus (Fig 2C). We found that 752 (51%) of the sites present in ProMetheusDB were already annotated, while the remaining 720 (49%) are novel. So, our repository significantly expands the knowledge on the extent of this PTM in human cells.

We compared mono-methyl-sites and di-methyl-sites to uncover a possible cross-talk between the two degrees of modification and observed that, of the total 1478 R sites, 783 are only mono-methylated, 290 are only di-methylated and 405 are identified in both forms (Fig 2D). To verify if these counts of mono- and di-methylated R residues reflect a bias in sample preparation procedure, we grouped our experiments based on whether they were based on the use of anti-pan-R-methyl antibodies from CST PTMScan kits for methyl-peptides enrichment or not. Indeed, the observation that mono- and di-methyl-R-sites were equally represented in the datasets deriving from experiments where no affinity-enrichment of R methylations was performed (Appendix Fig S2C) confirmed our hypothesis of a possible bias in the antibody specificity.

### Functional analysis of the R-methyl-proteome

To explore the potential impact of protein R methylation on cellular processes, we generated a network of functional protein:protein interactions (PPIs), starting from the full list of R-methyl-proteins included in ProMetheusDB. This analysis highlighted the presence of 8 sub-networks enriched for distinct biological pathways (Fig 3A and Appendix Fig S3A): the largest cluster consists of over 100 proteins and includes components of the spliceosome and other proteins involved in mRNA processing, confirming the well-known role of R methylation in RNA metabolism (Fong et al, 2019; Spadotto et al, 2020). The remaining 7 clusters included around 20-30 proteins each and were enriched for terms such as “DNA-binding transcription factor activity”, “Lipid metabolism”, “Cytoskeleton” and “Mitotic Anaphase and Metaphase”, “Fc Gamma receptor (FCGR)-dependent phagocytosis”, “Translation”, “Antigen processing” and “Chromatin organization” (Fig 3B).

**Figure 3.**
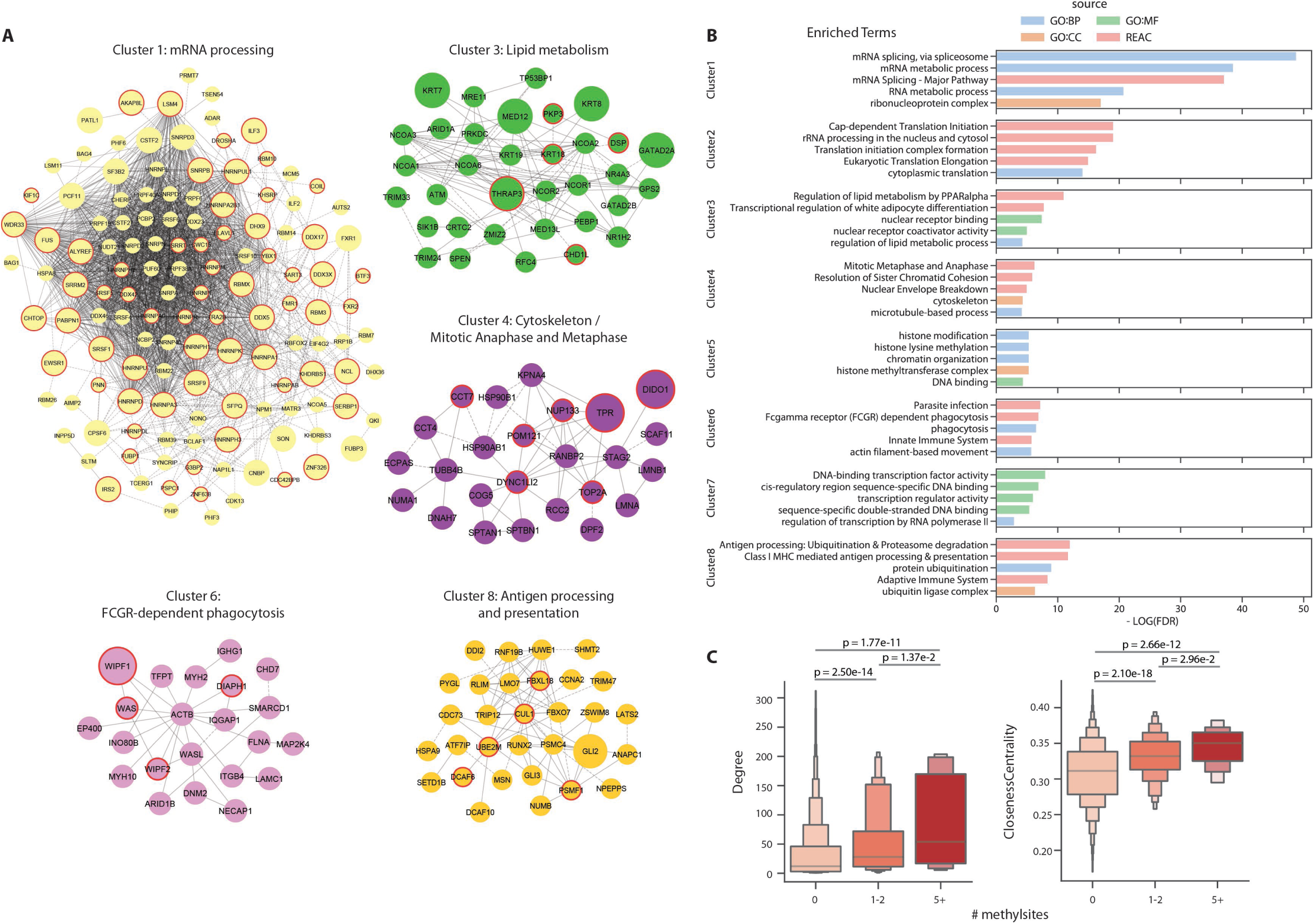
Functional analysis of the R-methyl-proteins network. **A)** Protein clusters identified in the Reactome functional interaction network of R-methyl-proteins. Larger nodes represent proteins with 5 or more methylation sites (hyper-methylated), while red borders highlight proteins bearing regulated methyl-sites. **B)** Top 5 most enriched functional GO terms found for each protein cluster (GO:BP = Gene ontology biological processes; GO:CC = gene ontology cellular component; GO:MF = Gene ontology molecular function; REAC = Reactome pathways). **C)** Topology analysis performed on the Reactome interaction network reveals an increase in degree and centrality of proteins at increasing numbers of R-methyl-sites (p-values calculated with Kruskal-Wallis test).

Since we previously observed a strong trend of methylated proteins to be subunit of complexes (Bremang et al, 2013), we asked whether this observation still held true in this expanded dataset: we found that hyper-methylated proteins (bearing five or more R methyl-sites) presented significantly higher node degree and network centrality, not only compared to non-methylated proteins (detected in the Input samples; p = 1.77e-11 and p = 2.66e-12) but also to hypo-methylated ones (presenting one or two R-methyl-sites; p = 0.01 and p = 0.03, respectively; Fig 3C). A similar analysis considering only physical PPIs confirmed this result (Appendix Fig S3B). Several of the high-centrality, hyper-methylated proteins are known RBPs, such as RBMX, NONO, various ribosomal proteins, nucleolin and HNRNPs (Geuens et al, 2016). However, the presence of R-methyl-proteins in cellular pathways related to lipid metabolism, cytoskeleton organization and phagocytosis was particularly interesting. Among these proteins we found G3BP1, which is on the one hand linked to stress granule formation and on the other hand has been shown to participate in DNA/RNA-sensing pathways implicated in the regulation of innate immunity (Liu et al, 2019; Reineke & Lloyd, 2015); the cytoskeletal protein actin (ACTB); the nuclear receptors NCOA2 and NCOA3, which are involved in metabolism, inflammation and adipocytes differentiation (Rollins et al, 2015); CUL1, a component of several E3 ubiquitin-protein ligase complexes (Michel & Xiong, 1998); SMARCA5, a helicase with nucleosome-remodelling activity that has a role in transcription, phosphorylation of H2AX, and maintenance of chromatin structures during DNA replication (Oppikofer et al, 2017). Taken together, these proteins and pathways suggest a role of R methylation in innate and adaptive immunity, in line with published evidence linking PRMT1 to macrophage differentiation and apoptosis, inflammation and cytokine production (Cho et al, 2018; Tikhanovich et al, 2017; Zhao et al, 2019). Moreover, although all the aforementioned proteins were already annotated as methylated, our analysis revealed several novel R-methyl-sites on RBMX (R383, R388), NONO (R142), SMARCA5 (R616), HSP90B1 (R51, R557) and ACTB (R206, R312).

As a further investigation, we integrated the network of protein interactions with the quantitative data on methylation dynamics obtained from the SILAC experiments, to identify which proteins present at least one R-methyl-site that is significantly regulated in at least one SILAC experiment (i.e., regulated in the same way in one pair of Forward and Reverse replicates). Proteins featuring one or more regulated R-sites (circled in red in Fig 3A) showed higher node degree and centrality in the network (p = 0.04 and p = 0.03, respectively; Fig 4A); we confirmed this result when we repeated the analysis on a network of physical interactions only (Appendix Fig S3C). Moreover, we observed that this protein group was also enriched for the GO terms “RNA binding”, “RNA splicing” and “nucleic acid transport”, while proteins not bearing regulated methylations were not enriched for any specific GO term (Fig 4B). This suggests that two categories of R-methyl-sites may exist: “dynamic” methyl-sites whose changes serve to modulate protein:RNA interactions, and “structural” or “constitutive” methyl-sites, which occur on a wider range of protein types, remain stable upon different stimuli and are likely linked to the life-cycle of the proteins they decorate.

**Figure 4.**
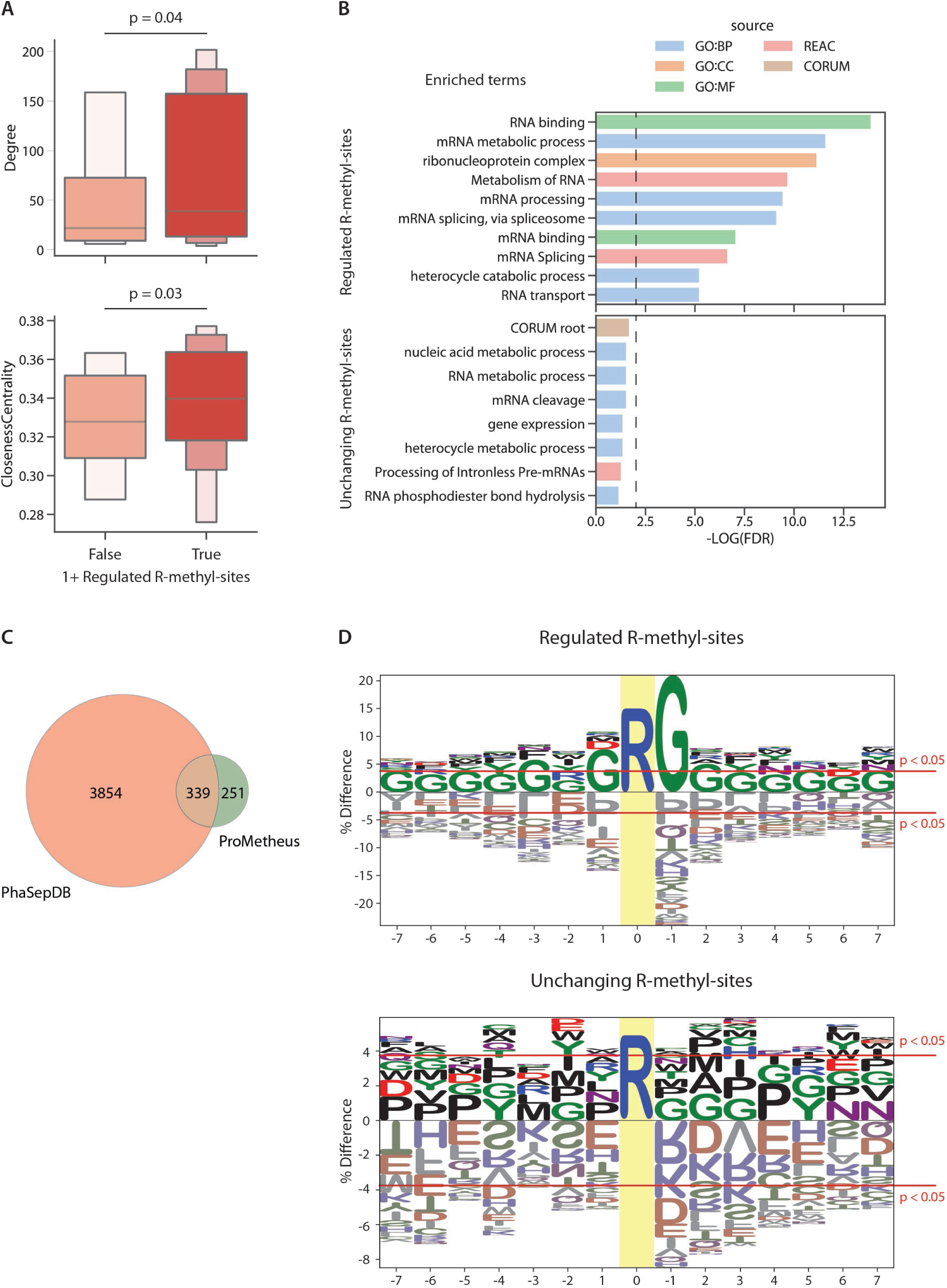
Functional analysis of dynamically regulated R methylations. **A)** Topology analysis performed on the Reactome network reveals that proteins bearing at least 1 significantly regulated R-methyl-site have significantly higher degree and network centrality than those without regulated R-methyl-sites (p-values calculated with Mann–Whitney test). **B)** Functional enrichment performed on proteins bearing at least one regulated R-methyl-site (top) or no regulated R-methyl-sites (bottom). (GO:BP = Gene ontology biological processes; GO:CC = gene ontology cellular component; GO:MF = Gene ontology molecular function; REAC = Reactome pathways; CORUM = CORUM protein complexes). **C)** Intersection of PhaSepDB and ProMetheusDB shows that 57% of R-methyl-proteins are involved in the process of liquid-liquid phase separation (LLPS). **D)** Motif analysis performed on significantly regulated (top) and unchanging (bottom) R-sites. Logos were generated using the full list of identified methyl-sites as background.

Since RNA binding proteins are frequently R-methylated and often involved in the process of Liquid-Liquid Phase Separation (LLPS), we asked whether a link could exist between R methylation and LLPS and intersected the ProMetheusDB with PhaSepDB, a database of proteins involved in LLPS: we found that >50% of the proteins in our dataset were also annotated as part of at least one membrane-less organelle (MLO) within PhaSepDB (Fig 4C). This observation, together with a very recent study published by our group (Maniaci et al, 2021), confirms that R methylation is involved in the assembly of MLOs.

Finally, because the major type I and II PRMTs (PRMT1 and PRMT5) preferentially target Rs within glycine/arginine-rich (GAR) motifs, we carried out a logo analysis on regulated R-methyl-sites versus the non-changing ones and found that regulated sites are surrounded by Glycines (Fig 4D, top). Instead, R-methyl-sites that emerged as unchanging in different functional states did not produce a significant enrichment for any consensus motif (Fig 4D, bottom): this result may suggest that unchanging methylations are deposed by methyltransferases other than the best-known ones that may recognize different sequence motifs, yet to be characterised.

### Structural survey of R-methylations hints at novel biological mechanisms

In the last two decades, intrinsically disordered regions (IDRs) of proteins have been reported to play a role in different processes (such as protein:protein and protein:nucleic acid interactions and protein phase separation) and often be modified by PTMs, including R methylation (Bremang et al, 2013; Darling & Uversky, 2018; Musselman & Kutateladze, 2021). In addition, methylated Rs can serve as docking sites for the recruitment of other proteins and the formation of multi-protein complexes; two examples of R methylation “readers” are represented by the Tudor domain, found in several proteins linked to transcriptional regulation, mRNA splicing and formation of RNP granules (Ying & Chen, 2012), and the WD40 repeat domain, located on a wide range of scaffolding proteins whose role is to bring together interactions partners into stable complexes (Jain & Pandey, 2018).

To study the link between protein structure and R methylation, we intersected the R-methyl-sites in the ProMetheusDB with disordered regions and domains annotated in MobiDB (Piovesan et al, 2021) and confirmed that the vast majority (77%) of the R-methyl-sites are indeed located in disordered regions (Fig 5A). When we carried out the same analysis on a subset of 1500 non-methylated R-sites randomly extracted from the human proteome, we observed a more even distribution between structured domains and disordered regions. As a matter of fact, when performing a Fisher exact test on the counts of methylated or unmodified R located in structured or disordered regions, we found that the distribution was not random (p = 5.15e-71; Fig 5A), which corroborates previous evidence that R-methyl-sites tend to occur much more frequently within low complexity and disordered regions.

**Figure 5.**
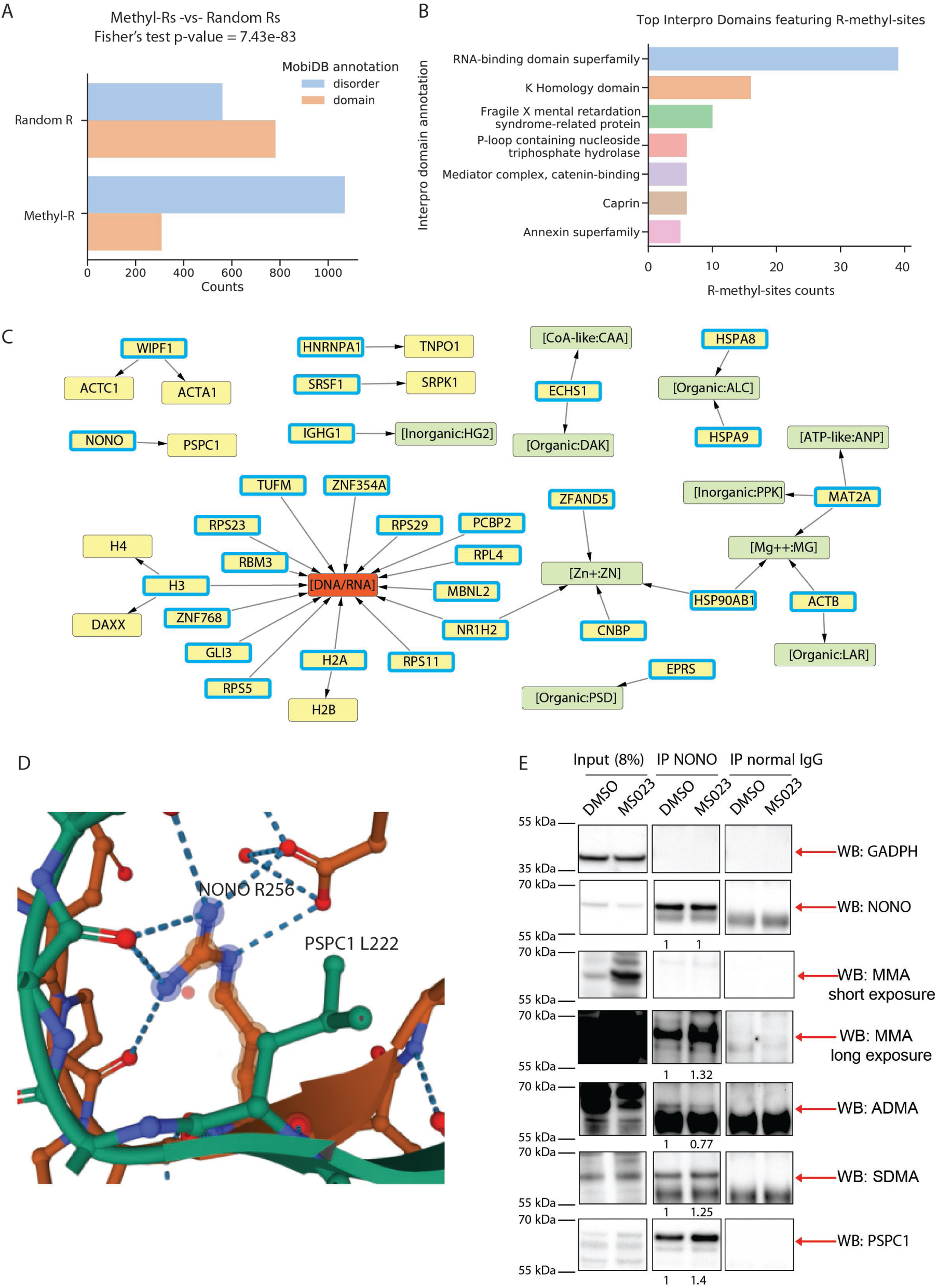
Structural analysis of R-methylated protein regions. **A)** Counts of R-methyl-sites that occur in regions that are annotated as either domains or disordered in the MobiDB database. Counts of Random R sites are also included as a comparison. P-value between R-methyl-sites counts and random sites counts was calculated with Fisher exact test. **B)** Counts of R-methyl-sites occurring on specific protein domains, as annotated in the InterPro database. Domains linked to RNA-binding proteins such as RNA binding domain and K homology domain are the most represented. **C)** Network obtained from Mechismo, showing the methylated proteins in ProMetheusDB and their interaction partners. DNA/RNA and chemicals are highlighted in orange and green, respectively. R-methylated proteins are indicated by a blue border. **D)** Crystal structure of NONO (orange) and PSPC1 (green) showing how R256 of NONO, which is annotated as methylated in ProMetheusDB, interacts with L222 of PSPC1, suggesting potential role in the binding of the two proteins. **E)** IP of NONO and Western blot (WB) analysis of NONO, its R methylation state and PSPC1 co-enrichment, in cells treated with either DMSO (negative control) or with the small molecule inhibitor MS023. Increased levels of NONO-MMA and, to a lesser extent, NONO-SDMA and the PSPC1 co-immunoprecipitated were measured upon MS023 treatment, compared to DMSO. Quantification of signal intensity for each band was performed by FiJi software and normalized as described in Material & Methods.

Despite being a minority, we focused on the 308 methyl-Rs located in structured regions and mapped them onto InterPro (Blum et al, 2021) domains to investigate their possible association to specific protein domains: most R-methyl-sites were found to localize within “RNA Binding Domain”, “K Homology domain” and on the “Fragile X mental retardation syndrome-related” family of proteins (Fig 5B). Aside from the “RNA Binding Domain”, whose enrichment is expected, the K Homology domain is also mainly located in RBPs (specifically heterogeneous nuclear ribonucleoproteins (Valverde et al, 2008)) and the Fragile X mental retardation syndrome-related proteins also belong to the RBP family; hence, these results are consistent with the notion that RNA binding and processing are highly dependent on this PTM.

We then set out to investigate the possible role of R methylation in regulating the interaction of proteins with other types of biomolecules. To do so, we analysed ProMetheusDB using Mechismo, a web application that maps alterations of amino acid residues (induced by mutations or PTMs) onto protein crystal structures, to identify modified sites that are putatively involved in physical interactions with different kinds of biomolecules (Betts et al, 2015; Fic et al, 2021). We found that 26 methylated Rs were involved in a total of 65 interactions with nucleic acid (n = 11), chemical compounds (n = 7), other copies of the same protein (n = 5) and other proteins (n = 42) (Fig 5C). The presence of more interactions with proteins rather than with nucleic acids for these modifications located in structured domains is somewhat unexpected, but it can be explained by the fact that many RBPs bind RNA through low complexity regions, for which crystal structures (on which Mechismo relies on) are currently not available. Still, these data support the hypothesis that R methylation can also modulate interactions with biomolecules other than RNAs, such as protein:protein and protein:chemical.

We followed up the R-methyl-sites indicated by Mechismo analysis as putatively involved in protein:protein interactions: based on reports that methylation of the guanidino-group of arginines can exert distinct effects on protein:protein interactions by reducing the number of hydrogen bond donors on this amino acid, while - at the same time-increasing its hydrophobicity (Fulton et al, 2019), we hypothesized that methylation could on the one hand enhance interactions between Rs and hydrophobic residues and on the other hand inhibit the interactions between Rs and negatively charged amino acids.

An interesting case-study highlighted by the structural analysis was represented by the protein-pair NONO:PSPC1, where R256 of NONO is located at the interface of interaction with PSPC1 (Fig 5D). We hypothesized that methylation of R256 would increase its hydrophobicity, thus stabilizing the NONO:PSPC1 interaction interface, which also entails a-polar residues, such as L222 of PSPC1. To verify this prediction, we set up the NONO immuno-precipitation (IP) in HeLa cells and profiled PSCP1 co-immunoprecipitation efficiency, both in basal conditions and upon treatment with the PRMT type I inhibitor MS023 (Eram et al, 2016), which typically leads to ADMA reduction and MMA/SDMA increase, depending on the enzyme processivity on a specific site and/or the scavenging effects by other PRMTs, as reported in (Dhar et al, 2013; Hartel et al, 2019; Musiani et al, 2020). When we profiled both NONO R methylation state and interaction with PSPC1, we observed that MS023 induced the decrease of ADMA on NONO, with a parallel increase of MMA and, to a minor extent, of SDMA; in parallel, a mild increase of the amount of PSPC1 co-immunoprecipitated was also observed, which confirms our prediction (Fig 5E).

### Cross-talk between R methylation and S/T-Y phosphorylation

The cross-talk between R methylation and S/T-Y phosphorylation has already been described in the literature and linked to subcellular localization of proteins (Smith et al, 2020) and the promotion of stem-like properties in cancer (Liu et al, 2020). Interestingly, phosphorylation and methylation share some features like the preferential localisation in IDRs, the role in modulating protein:RNA interactions and LLPS (Hamey et al, 2021; Owen & Shewmaker, 2019; Schisa & Elaswad, 2021). To study this aspect more in-depth, we determined 15-amino acid windows centred on each R-methy-site annotated in our ProMetheusDB and assessed if phosphorylated residues (see Methods) are preferentially enriched within these sequence windows. We found that 706 (47.8%) of the 1478 R-methyl-sites in our dataset present an S, T or Y phosphorylation site within their 15-amino acid window; by repeating this analysis on 1500 R residues randomly selected from the human proteome, we observed that the percentage of sites featuring a phosphorylation site in the same 15-amino acid window was halved (356 sites, corresponding to 23.7%; Fig 6A), which indicates a statistically significant overrepresentation of phosphosites in the proximity of methyl-Rs (p = 4.62e-43), thus supporting the hypothesis of the co-occurrence of these PTMs.

**Figure 6.**
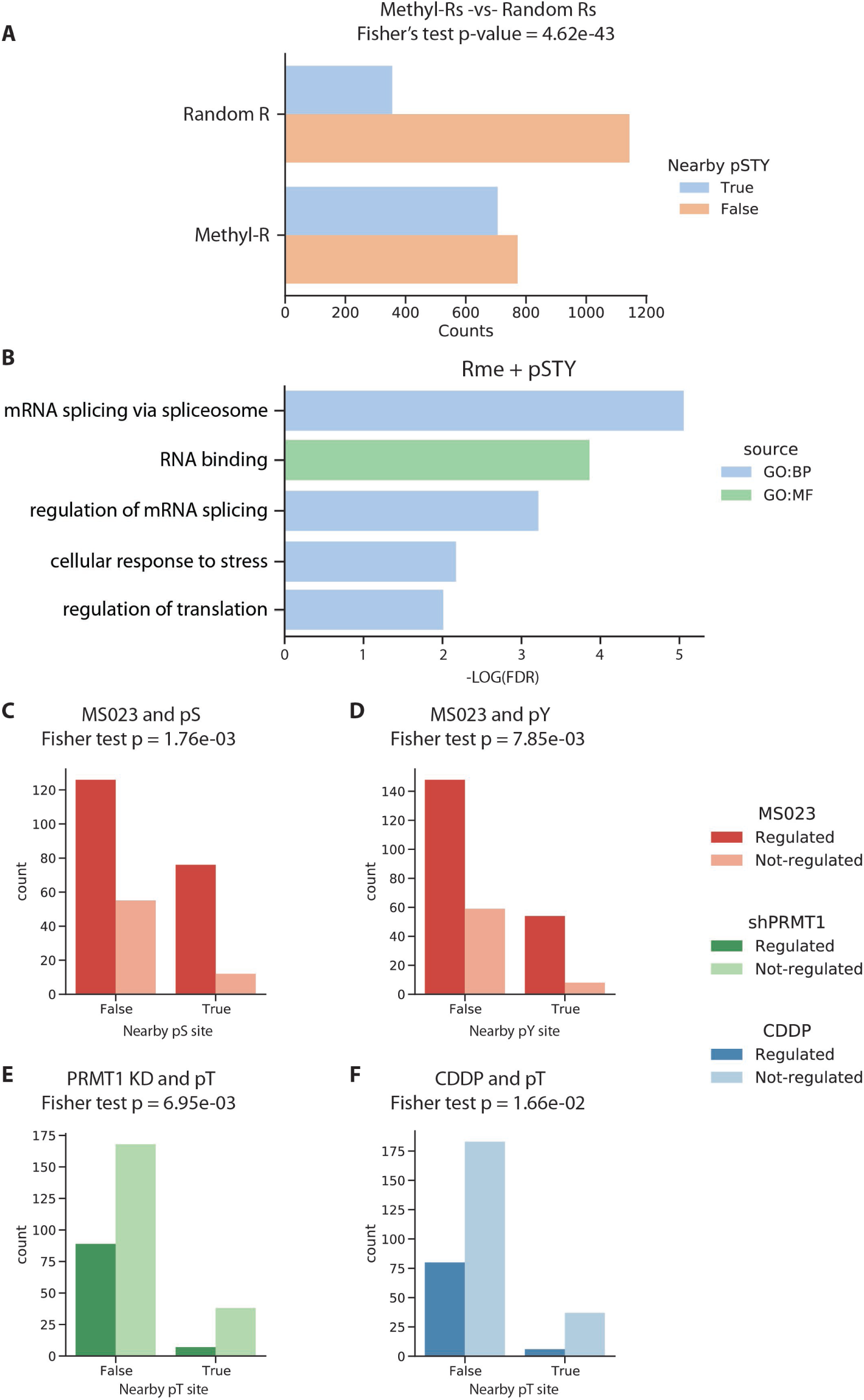
Cross-talk of R methylation and S/T-Y phosphorylation. **A)** Counts of R-methyl-sites that occur in proximity of a phosphorylated site. As a control, counts of randomly sampled R sites from the human proteome are shown, and the comparison indicates a significant correlation between methylation and phosphorylation (p-values calculated by Fisher’s exact test). **B)** Functional terms enriched from proteins bearing proximal R methylation and S/T-Y phosphorylation sites (GO:BP = Gene ontology biological processes; GO:CC = gene ontology cellular component; GO:MF = Gene ontology molecular function; REAC = Reactome pathways; KEGG = KEGG pathways; CORUM = CORUM protein complexes). **C-F)** Counts of R-methyl-sites classified based on i) their regulation state upon external stimuli and ii) their proximity to S/T-Y phosphorylation sites (p-values calculated by Fisher’s exact test).

We then asked whether the co-occurrence of methylation and phosphorylation was linked to a specific biological process by performing GO analysis on the protein displaying a statistically significant co-occurrence of R-methyl-sites and phospho-STY sites: we found that these proteins are even more strongly linked to RNA binding and splicing compared to other proteins in ProMetheusDB (Fig 6B). To assess whether R-methyl-sites co-occurring with phosphosites are more likely subject to regulation in response to external cues, we intersected the results of the PTM cross-talk analysis with the dynamic information derived from the SILAC experiments, where methylation changes have been profiled in response to PRMTs type I inhibition by MS023, cisplatin (CDDP) treatment and PRMT1 knock-down (KD) or overexpression (OE). Despite being very different stimuli, we expected to observe an effect on PRMT1 methylation targets in all cases, as it is the most active type I enzyme (and thus more affected by MS023 than other PRMTs) and CDDP treatment causes re-localization of PRMT1 to chromatin (therefore inducing a decrease in the methylation levels of cytosolic proteins). Interestingly, we found that phospho-S and phospho-Y sites seem to occur more frequently in the proximity of the MS023-regulated methyl-sites than the non-regulated ones (p = 1.76e-03 and p = 7.85e-03; Figure 6C-D). Moreover, the R-methyl-sites unchanging upon either CDDP treatment or PRMT1 KD tend to present a phospho-T site in their surrounding 15-amino acid sequence window (p = 6.95e-03 and p = 1.66e-02; Figure 6E-F). The different responses to these stimuli suggest that there are multiple ways through which these PTMs can influence each other, which hints towards the widespread existence of a molecular barcode of PTMs, similar to that extensively described on histones

We then expanded the analysis of possible cross-talk of R methylation with other PTMs, such as K acetylation, ubiquitination and sumoylation and found that R-methyl-sites seem to significantly anti-correlate with nearby K ubiquitination sites, when compared to the dataset of randomly selected R-sites (Appendix Fig S4A). Instead, the associations between R methylation and K acetylation or sumoylation are not statistically significant (Appendix Fig S4B-C). Because the numbers of acetylation, ubiquitination and sumoylation sites annotated in Phosphosite Plus are at least one order of magnitude lower than that of phosphosites and this difference might introduce a bias in the analysis, these results should be taken with caution; nevertheless, they seem to corroborate the specificity of cross-talk between R methylation and S/T-Y phosphorylation, which is not generically applicable any annotated PTM.

### Methylation beyond Arginine: hmSILAC-based detection of non-canonical methylation sites

Protein methylation has been observed not only on Ks and Rs but also on residues such as Aspartate (D), Glutamate (E), Asparagine (N), Glutamine (Q), Serine (S) threonine (T) and Histidine (H). However, the systematic study on these non-canonical methylated residues by MS is hindered not only by the high FDR that already plagues R methylation studies but also by the difficulty in pinpointing the PTM to its exact residue when multiple putative methyl-sites are present on the same peptide. In this context, we set to assess to what extent the hmSILAC strategy could help filling this gap in knowledge. With the aim of exploiting at their maximum potential the hmSILAC strategy and the large set of MS data available, we re-analysed all the raw data from hmSILAC-labelled, non-affinity enriched samples with the Andromeda search engine of MaxQuant, allowing mono-methylation (either light or heavy) to occur not only on R and K but also on D, E, N, Q, S, T and H. Our rationale was that all enzymatically-driven methylations of both standard and non-conventional residues should use SAM as the universal methyl-group donor.

MaxQuant output data were then processed with hmSEEKER v2.0 using the same criteria employed for the assignment of the high-quality R-methyl-sites (i.e. peptide score > 25; PTM localization probability > 0.75; use of ML model to identify true doublets) to produce a list of orthogonally validated, methylation sites which included 111 methylation sites in total, 42 of which occurring on non-canonical residues located on 30 different proteins (Fig 7A). These novel modifications are mainly found on proteins functionally linked to mRNA splicing and RNPs (Fig 7B). Although most proteins only present one type of methylated residue, combinations of up to 5 different methyl-residues appeared on a few proteins (Fig 7C), such as TAF15 (experimentally found to be methylated on R, K, S, D and Q), HSPA8 (methylated on K, D, E and N) and EEF1A1 (methylated on K, S, D, Q). The methylated residues that co-occur most frequently seem to be R and S, which coexisted in five proteins (HNRNPD, HNRNPK, SFPQ, TAF15 and VIRMA). All the 46 R-methyl-sites and 20 K-methyl-sites that had been previously identified in the search for R/K methylations were re-identified in this search; 3 K-methyl-sites unique for this second search were identified in combination with other non-canonical methyl-sites, therefore Andromeda could not identify them during the first search, that was constrained to K an R methylation only.

**Figure 7.**
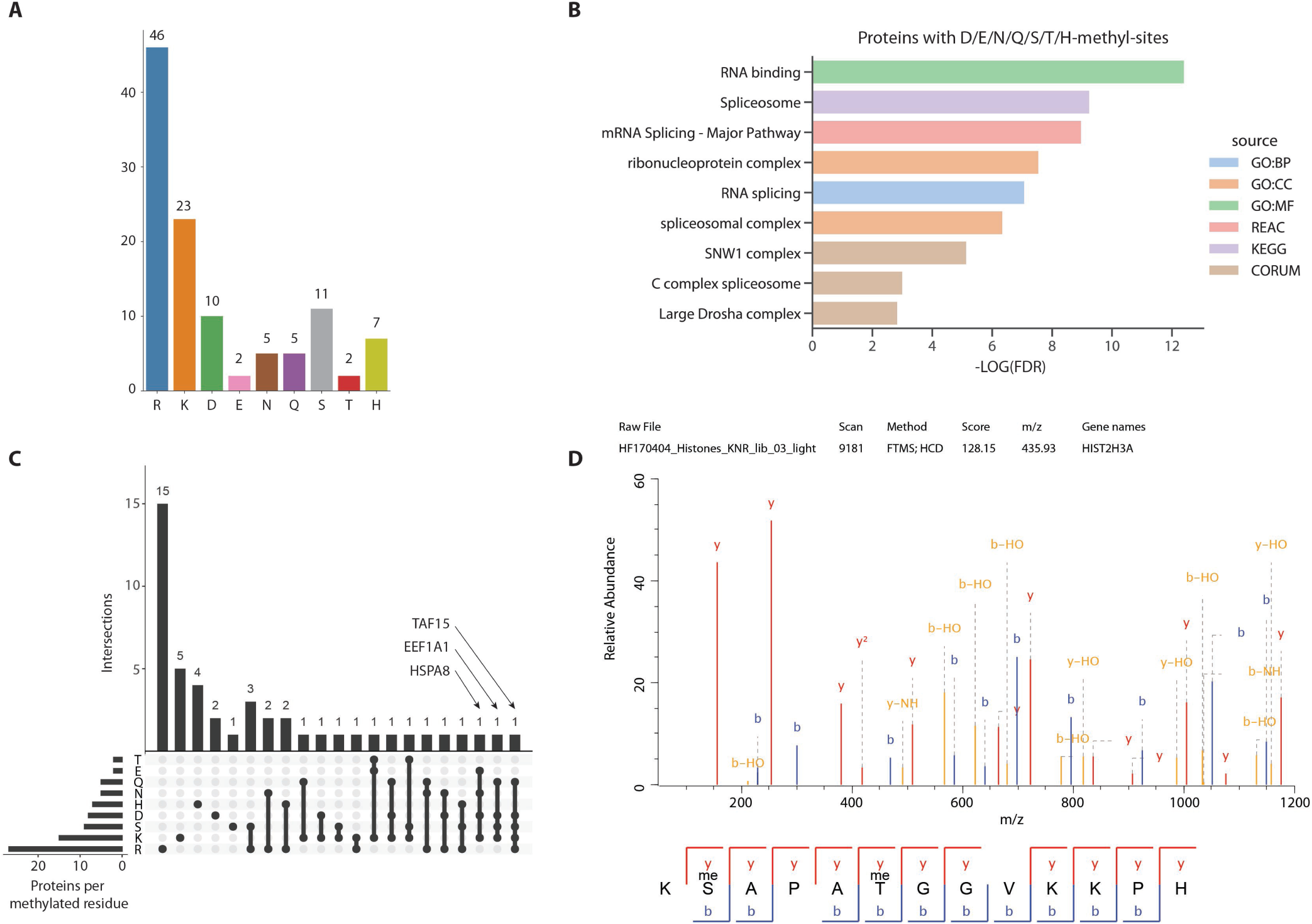
Explorative analysis of non-canonical methylation sites. **A)** Number of methyl-sites detected upon the re-analysis of the Input MS data, grouped by residue. **B)** Gene Ontology terms, pathways and complexes significantly enriched among proteins bearing methylations on non-K/R residues. **C)** UpSets plot representation of the co-occurrence of methylated residues on different proteins. **D)** MS/MS spectrum of the histone H3 27-40 peptide methylated on S28 and T32.

This exploratory analysis also revealed the presence of two unconventional novel methyl-sites on H3, on S28 (H3S28me) and T32 (H3T32me). We were thus prompted to focus on non-canonical methylation on histones by this initial evidence, together with two remarks: first, histone methylation is widely regarded as a core component of the histone code, hence the detection of novel methyl-sites may help dissecting this molecular language affecting gene expression; second, since standard database search engines produce suboptimal results when multiple (>5) PTMs are searched in a large protein database (Verheggen et al, 2020), limiting the analysis to histones could be a reasonable trade-off to expand our methylation search to non-canonical methyl-residues while circumventing the explosion of false positives and negatives linked to the expansion of search space.

We thus applied an optimized biochemical and analytical pipeline to a set of hmSILAC-labelled histone samples for the in-depth identification of methylation sites on these proteins; from a biochemical point of view, we took advantage of multiple proteases (i.e. Trypsin, LysC, ArgC and LysargiNase) digestion of histones to generate overlapping peptides and maximise protein coverage. From the computational point of view, the MS raw data were searched with MaxQuant using a filtered version of the UniProt Human protein database that included only histone proteins, using the same search parameters selected for the global methylation search. The subsequent analysis with hmSEEKER v2.0 for methyl-peptides pair matching and orthogonal validation of novel methyl-sites allowed us to unambiguously re-identify mono-methylation on H3S28 and H3T32 (Fig 7D; Appendix Fig S5).

Follow up studies relying on the production and use of ChIP-seq grade antibodies as well as synthetic biotinylated peptides for screening of putative interactors will be the gateway to retrieve experimental data to hypothesize the functional role of these novel histone PTMs and their cross-talk with already known ones.

## DISCUSSION

In this work, we describe ProMetheusDB, our current hmSILAC-validated repository of protein methylation sites, annotated upon the re-analysis of previous hmSILAC experiments with an updated version of the hmSEEKER bioinformatics tool. Doublets of light and heavy methyl-peptides were evaluated by a ML model that was trained on doublets generated by M-containing peptides, which, unlike proper methyl-peptides, can be easily identified by database search engines and therefore provide a reliable positive control to our tool. After training the model, we observed that the dRT parameter of the doublets was the least important for discriminating true and false peptide pairs; this is in contrast with the first iteration of hmSEEKER, where dRT and ME were the main predictors of methyl-peptide doublets and the H/L ratio parameter was introduced later. This choice of features also differentiates our ML model from the one adopted by MethylQuant (Tay et al, 2018), which scores methyl-peptide pairs based on the isotope distribution and elution profile correlation of the two peaks.

This study focused on R methylation, which over the last decade has been found to be a key player in a variety of biological processes. The functional analysis performed on R-methyl-sites revealed that proteins carrying either multiple R-methyl-sites and/or dynamically regulated ones are more strongly interconnected with other proteins in our Reactome-derived network, thus representing potential hubs of protein:protein interactions.

Interestingly, the protein clusters emerging from the network analysis suggest a potential role of R methylation in the immune response, as exemplified by the “FCGR-dependent phagocytosis” cluster. While it is known that PRMT1-mediated H4R3me2a promotes the expression of PPARgamma, a transcription factor regulating monocyte differentiation into anti-inflammatory macrophages (Tikhanovich et al, 2017), our results expand the role of R methylation in immunity beyond mere transcriptional regulation, showing that several methyl-proteins are involved in antigen processing and exogenous DNA/RNA sensing pathways (such as CUL1 and G3BP1, respectively). Other clusters that might be connected to this biological process are the ones related to cytoskeleton dynamics and lipid metabolism. Anti-inflammatory macrophages metabolize lipids as a source of energy, while inflammatory ones use fatty acids to produce prostaglandins and leukotrienes, which act as hormone-like signal molecules to regulate the inflammatory response (Batista-Gonzalez et al, 2019; Yan & Horng, 2020). The impact of R methylation on proteins of the cytoskeleton (such as actin) has been already described in neuronal development, where PRMTs regulate the formation of both the axon and the dendrites; it is possible PRMTs control cytoskeleton dynamics also in macrophages and potentially other cell types.

By mapping the R-methyl-sites annotated in ProMetheusDB onto protein structures, we confirmed that this PTM mostly occurs in unstructured, low-complexity regions, which are involved in mediating protein:RNA interactions. Intrinsically disordered regions are characterized by short linear motifs well-established as important mediators of protein:protein interactions (PPIs) (Tompa et al, 2014). Our observation that R methylation most frequently occurs in IDRs, together with the notion that heavily R methylated proteins are also central nodes within functional PPI network, is indicative of a functional link between R methylation and PPI modulation. Indeed, by inspecting available 3D complexes, we also found a small number of sites where R methylation could modulate protein:protein interactions by increasing or reducing the hydrophobicity and hydrogen-bonding capability of R situated at protein interfaces.

Our data and a recent study by Xiang-Bo Wan and co-workers (Yin et al, 2021) identify NONO R256 and R251, respectively, as methylation sites. Both residues are predicted by Mechismo to interact with a-polar residues of PSPC1 (L171 and L222, respectively). Moreover, the amino acid substitution of NONO R256 with an Isoleucine (I) is a mutation associated with colorectal cancer in the ActiveDriverDB database (Krassowski et al, 2018) and is also predicted by Mechismo to enhance the NONO:PSPC1 interaction; similarly, a hypothetical R251I substitution is also predicted to strengthen the binding. Overall, these observations suggest that PRMT1 may elicit an oncogenic effect (at least in colorectal cancer) by modulating the interaction between NONO and PSPC1 through the methylation of NONO R251 and/or R256.

By contrast, we hypothesize that methylation of SRSF1 on R154 might inhibit the interaction between SRSF1 and SRPK1. SRPK1-mediated phosphorylation of SRSF1 regulates alternative splicing and promotes the expression of protein variants that have anti-apoptotic and pro-angiogenic properties (Nowak et al, 2008). In this context, Jacky Chi Ki Ngo and collaborators showed that blocking the interaction of SRPK1 and SRSF1 with a PPI inhibitor could reduce SRSF1 phosphorylation and thus suppress angiogenesis (Li et al, 2021). Our data suggest that a similar result could be achieved by regulating SRSF1 methylation levels, although more in-depth mechanistic studies are needed to confirm this hypothesis.

An additional interesting interaction suggested by Mechismo involves the R264 of SAM Synthase (MAT2A), which is located at the interface between the subunits that form the active dimer of the enzyme, where it contacts E57, A281 and K285 of the other MAT2A monomer. This residue also interacts with the substrate ATP and the metal ions serving as cofactors for the reaction. The observation that the enzyme that synthesizes SAM, the methyl-group donor, can also be methylated in this relevant pocket for its catalytic activity is particularly intriguing, because it may suggest the existence of a feedback loop controlling SAM levels in the cell. It has already been reported that the RNA methyltransferase METTL16 binds and methylates the 3’-UTR region of MAT2A mRNA to prevent its translation. When SAM levels are low, the mRNA of MAT2A cannot be methylated due to lack of the methyl donor and the protein is translated (Pendleton et al, 2017). MAT2A protein methylation at R264 could represent an additional layer of post-translational regulation of the enzyme.

Our structural analysis, however, suffers from two limitations: first, methylation often occurs in disordered regions which cannot be studied through classical crystallography approaches; second, not all the structures of proteins that can be crystallized have necessarily been solved. However, thanks to the advancements in the field of AI, the scientific community now has access to AlphaFold, which can predict a protein structure from its primary sequence with unprecedented speed and accuracy (Tunyasuvunakool *et al*., 2021). Therefore, as a future perspective, it could be interesting to expand our analysis using AI-predicted protein structures.

Phosphorylation and methylation sites co-localize on disordered SRGG motifs of some proteins, where the two PTMs are mutually exclusive because the presence of negative charges inhibits the binding of PRMTs to their recognition motifs (Smith et al, 2020). By overlapping ProMetheusDB with the Phosphosite Plus phosphoproteomics dataset, we confirmed that methylated Rs are significantly more likely to occur in the proximity of a phospho-S but also expand this observation to phospho-T and -Y residues. Furthermore, the analysis of co-occurrence of these two PTMs in the context of dynamic regulation suggests the existence of a link between the regulation of an R-methyl-site and its proximity to a phosphosite. Acute stimuli such as treatment with MS023 (which inhibits type I PRMTs, with a preference for PRMT1 at the experimental condition used in our studies) cause a change in the methylation of Rs that are close to phosphosites; instead, modulation of PRMT1 expression and the treatment with the chemotherapeutic drug cisplatin affect methyl-Rs that are distant from phosphorylation sites. This result highlights the importance, in the future, of developing experimental pipelines enabling the simultaneous profiling of these two (or even more) PTMs, something that is currently still at a pioneering stage (Hamey et al, 2021). Our analysis, for instance, focused exclusively on methyl-sites, whereas the datasets of phosphorylation, acetylation, ubiquitination and sumoylation sites were mostly derived from other studies focused on one individual PTM at a time. Profiling differently modified isoforms of a peptide (e.g. unmodified, methylated, phosphorylated, co-modified) within a single experiment would be very informative to experimentally assess which PTMs can truly co-exist and which are mutually exclusive, both a basal state but also during transition or in response to external cues.

The application of the hmSILAC biochemical and analytical pipeline led to the annotation of methyl-sites also on amino acids that are traditionally excluded from proteome with MS-based analysis, such as D, E, N, Q, S, T and H, which also included the identification of a few putative novel methyl-marks on histones (i.e., H3S28, H3T32). The two PTMs occurring on the peptide 27-40 of histone H3 are particularly interesting for their potential cross-talk with functionally relevant modifications on K27 and K36. For instance, methylation of H3K27 is a repressive mark (Wiles & Selker, 2017), whereas methylation of H3K36 plays a role in transcriptional regulation, whereby H3K36me2 counteracts gene silencing by blocking the recruitment of PRC2 complexes; however, when genes are transcribed, this mark is replaced by H3K36me3 to prevent transcription initiation from intragenic regions (Huang & Zhu, 2018). Our experimental data showed that the S28me mark could coexist with K27me and T32me, while peptides carrying simultaneously K27me and T32me, or bearing S28me/T32me in combination with K36me were not detected (Appendix Fig S5). A more in-depth analysis of this peptide and its numerous differentially modified isoforms is needed to corroborate this observation: measuring the relative abundance of the K27me/S28me and S28me/T32me peptide isoforms could allow developing some hypotheses on their biological role and cross-talk with other epigenetic marks. Similarly, it would be interesting to confirm whether S28me/T32me are mutually exclusive with K36 methylation or to find how these novel methyl marks are associated with neighbouring acetylation and phosphorylation.

We argue that our analysis of non-canonical methylation sites could be linked to our previous analysis on PTMs cross-talk. A recent paper from the Eyers’ group (Hardman & Eyers, 2020) explored the field of non-canonical phosphorylations and identified several phosphosites on R, K, D, E, H and C residues. Interestingly, these residues overlap with the putative non-canonical methyl-residues we have investigated here: it is, therefore, possible to envisage that the cross-talk of methylation and phosphorylation is not limited to proximal sites, but that may also occur through the physical competition for the same substrates/residues. Two possible case-studies emerging here are H3S28 and R55 of Keratin type I cytoskeletal 18 (KRT18R55). H3S28 is a known phosphorylation site that we identify as methylated in our hmSILAC histone dataset. Phosphorylation of H3S28 can occur in response to extracellular stress and is proposed to override repressive epigenetic marks to temporarily express genes that would normally be silenced (Sawicka & Seiser, 2012). A reasonable hypothesis is that methylation of H3S28 may cooperate with methylation of H3K27 in silencing genes (Wiles & Selker, 2017), which would explain why we did not detect H3S28 in combination with the aforementioned active transcription marks H3K36me2/me3 and H3K27ac (Zhang et al, 2020). As a future perspective, since H3S28ph is also necessary for chromatin condensation (Sawicka et al, 2014), we could investigate how this residue is modified specifically during mitosis.

KRT18R55 is a methyl-site that was first identified in (Guo et al, 2014) and orthogonally validated in our hmSILAC experiments. The site is also present in the list of non-canonical phosphosites published by Eyers and co-workers, thus could be another example of competition between PTMs. Unfortunately, there is not enough data in the literature to make a hypothesis on the function of this modification site.

Some major points remain to be addressed in the methyl-proteomics field. First, there is a need for more efficient workflows that allow to annotate R-methyl-proteomes through sensibly smaller scale experiments, as the prerequisite for the investigation of this PTM in more relevant model systems, such as primary cells, tissues, organoid, similarly to what was made possible in the phospho-proteomics field (Lindhorst & Hummon, 2020). The implementation of Data-Independent Acquisition (DIA) methods could also prove beneficial by reducing the identification bias towards the most abundant proteins and thus increase the depth of coverage of the methyl-proteome (Bekker-Jensen et al, 2020).

Better strategies for the enrichment of di-methyl-R-peptides are also needed to address the bias introduced by the use of the CST anti-MMA kit. Although quantification of intracellular methyl-Rs derived from degradation of methylated proteins showed that ADMA is the most abundant of the three (Davids & Teerlink, 2013), most MS-based methyl-proteomics studies have so far reported a significantly larger number of MMA sites than ADMA/SDMA sites. We already discussed this issue, suggesting that the anti-pan-MMA antibodies traditionally used to enrich methyl-peptides outperform anti-pan-ADMA and SDMA ones thus generating a bias in methyl-proteomics studies (Spadotto et al, 2020). ProMetheusDB also suffers from this limitation, as highlighted by the fact that mono- and di-methylated R-sites are more evenly represented in the datasets deriving from experiments when no affinity-enrichment steps through the CST kits were performed (e.g. fractionated whole-cell extract, protein IPs; Appendix Fig S2C). Along the same line, there is a strong need for efficient antibodies for the enrichment of K-methyl-peptides to map the non-histone K-methyl-proteome. The availability of such reagents would allow researchers to finally address the question whether K methylation is widespread beyond histone, similarly to K acetylation and R methylation, regulating cellular processes beyond chromatin-based transcriptional regulation.

Besides the annotation of the steady-state methyl-proteome, we recognize a need to accelerate the acquisition of dynamic data. The triple SILAC strategy we adopted for some experiments is overall laborious and limited in its multiplexing capabilities. In the near future it would be useful to assess the feasibility of applying isobaric mass tags (TMT) to MS-based methyl-proteomics profiling, to overcome this limitation and perform multiplexed experiments where methylation changes can be profiled across multiple experimental conditions.

From the analytical point of view, some future developments can also be conceived: first, spectral library searching (Lam, 2011) is faster than conventional database searching (which can take several weeks when non-canonical methyl-sites are considered), however its application to methyl-proteomics was limited by the low number of high-confidence methyl-peptides spectra available. Within this context, the MS/MS spectra of methyl-peptides that were orthogonally validated by hmSILAC could be used to build a spectral library to be used as reference, thus speeding up the analysis of new data. Second, it is crucial to separately analyse ADMA and SDMA, which can be distinguished by searching for their diagnostic neutral loss ions within the MS/MS spectra (Brame et al, 2004; Musiani et al, 2019). However, at the moment, a systematic method for the automatic annotation of these characteristic neutral losses is still lacking; while general purpose tools can be adapted to perform this task (Kelstrup et al, 2014), the aforementioned lack of high-confidence spectra makes this task difficult. For these reasons, it is still recommended to visually inspect the MS/MS spectra of di-methyl-R-peptides, which is impractical in the case of large datasets such as ProMetheusDB.

## Materials & Methods

### Preparation of samples for global R methylation analysis

The annotation of ProMetheusDB was obtained by combining datasets derived from two distinct proteomics strategies. The majority of the methylations were annotated from hmSILAC experiments that allow this PTM to be orthogonally validated, as described in (Ong & Mann, 2006). This initial list was then integrated with a second list of regulated methyl-peptides derived from MS-proteomics experiments where the biochemical protocol to separate and affinity-enrich methyl-peptides was coupled to metabolic labelling with standard SILAC amino acids (i.e., K and R) and stimulation of cells with different perturbations (e.g. from pharmacological inhibition of genetic modulation of PRMTs, to treatment with genotoxic drugs). Each SILAC experiment was done in duplicates (forward and reverse) and only peptides whose regulation was reproducible in at least one pair of experiments was considered significantly regulated. As such, the experiments we analysed to build the methyl-proteome presented in this study are described in Table EV1.

### Extraction and protease digestion of hmSILAC-labelled histones before LC-MS analysis

hmSILAC labelled HeLa S3 cells (L and H channels mixed in 1:1 ratio) were resuspended in lysis buffer (10% sucrose, 0.5mM EGTA, 60mM KCl, 15mM NaCl, 15mM HEPES, 0.5mM PMSF, 5μg/ml Aprotonin, 5μg/ml Leupeptin, 1mM DTT, 5mM NaButyrate, 5mM NaF, 30μg/ml Spermine, 30μg/ml Spermidine and 0.5% Triton X-100) and nuclei were separated from cytoplasm by centrifugation on sucrose cushions for 30 minutes at 3,695 g, 4°C. Histones were then extracted through 0.4M hydrochloric acid for 5 hours at 4°C and dialyzed overnight in CH3COOH 100 mM Dialysed histones were then lyophilized and either kept at −80 until use or directly resuspended in milliQ water before sample processing prior to MS analysis. To maximize the protein sequence coverage, four different aliquots of 5 μg histones each were in-solution digested overnight using different proteases, such as Arg-C, Trypsin, LysargiNase and LysC. Proteolytic peptides were then desalted and concentrated by micro-chromatography onto SCX and C18 Stage Tips micro-column, prior to LC-MS/MS analysis.

### LC-MS/MS analysis

Peptide mixtures were analysed by online nano-flow liquid chromatography-tandem mass spectrometry using an EASY-nLC 1000 (Thermo Fisher Scientific) connected to a Q Exactive instrument (Thermo Fisher Scientific) through a nano-electrospray ion source. The nano-LC system was operated in one column set-up with a 50cm analytical column (75 μm inner diameter) packed with C18 resin (easySpray PEPMAP RSLC C18 2M 50cm x 75 M, Fisher Scientific) configuration. Solvent A was 0.1% formic acid (FA) and solvent B was 0.1% FA in 80% ACN. Samples were injected in an aqueous 0.1% TFA solution at a flow rate of 500 nL/min. Peptides were separated with a gradient of 5–40% solvent B over 90 min followed by a gradient of 40–60% for 10 min and 60–80% over 5 min at a flow rate of 250 nL/min in the EASY-nLC 1000 system. The Q-Exactive was operated in the data-dependent acquisition (DDA) mode to automatically switch between full scan MS and MS/MS acquisition. Survey full scan MS spectra (from m/z 300-1150) were analysed in the Orbitrap detector with resolution R=35,000 at m/z 400. The ten most intense peptide ions with charge states ≥2 were sequentially isolated to a target value of 3e6 and fragmented by Higher Energy Collision Dissociation (HCD) with a normalized collision energy setting of 25%. The maximum allowed ion accumulation times were 20 ms for full scans and 50 ms for MS/MS and the target value for MS/MS was set to 10^6^. The dynamic exclusion time was set to 20s.

### MS raw data processing with MaxQuant

MS raw data were analysed using the integrated MaxQuant software v1.3.0.5 or v1.5.2.8 (for SILAC) or v1.6.2.10 (for hmSILAC), using the Andromeda search engine (Cox & Mann, 2008; Cox et al, 2011). In all MaxQuant searches, the estimated FDR of all peptide identifications was set to a maximum of 1%. The main search was performed with a mass tolerance of 4.5 ppm. A maximum of 3 missed cleavages was permitted, and the minimum peptide length was fixed at 6 amino acids. The June 2020 version of the Uniprot reference proteome (Proteome ID: UP000005640) was used for peptide identification. Other parameters were different, according to the experiment and the sample to be analysed.

- <u>For the hmSILAC experiments</u>, each MS raw data was analysed twice to identify Heavy and Light methyl-peptides separately. In the “Light” analysis, mono-methylation of K/R (+14.016 Da), di-methylation of K/R (+28.031 Da), tri-methylation of K (+42.047 Da) and oxidation of M (+15.995 Da) were specified as variable modifications; Carbamidomethylation of Cys (+57.021 Da) was indicated as fixed modification. In the “Heavy” analysis, heavy mono-methylation of K/R (+18.038 Da), heavy di-methylation of K/R (+36.076 Da), heavy tri-methylation of K (+54.114 Da) and oxidation of M were specified as variable modifications; Carbamydomethylation of Cysteine (C) and isotope-labelled Methionine (MS) (+4.022 Da) were indicated as fixed modifications. Enzyme specificity was set to either Trypsin/P or LysargiNase.
- <u>For the standard SILAC methyl-proteomics experiments</u>, we indicated K8+R10 and/or K4+R6 as SILAC labels. N-terminal acetylation (+42.010 Da), M oxidation, mono-methyl-K/R and di-methyl-K/R were set as variable modifications; Carbamidomethylation of C was set as fixed modification. Enzyme specificity was set to Trypsin/P.
- <u>For histone hmSILAC experiments</u>, MS raw data were also analysed twice. Mono-methylation (light or heavy) was allowed on K, R, D, E, H, Q, N, S and T and we included K acetylation (+42.011 Da) as a variable modification to address the fact that this modification is very abundant and likely coexists with methylation on histones. Enzyme specificity was set to Trypsin/P, ArgC, LysC or LysargiNase. Finally, to reduce search complexity, a database containing only human histone sequences was used.

### Validation of methyl-peptides with hmSEEKER 2.0

Identification of light and heavy methyl-peptides doublets was carried out with hmSEEKER (Massignani et al, 2019). Methyl-peptides identified by MaxQuant were filtered to only retain peptides with Andromeda Score >25, Delta Score >12 and Localization Probability of the methylation sites >0.75. hmSILAC doublets were reconstructed with hmSEEKER. Initially, hmSILAC 1.0 called a methyl-peptide doublet when two peaks had the same charge, |ME| < 2 ppm, |dRT| < 0.5 min and |LogRatio| < 1. The Machine Learning model developed in this study within hmSEEKER v2.0 was trained using Python package Scikit-learn v0.23.1, as described in the Results section.

### Downstream quantitative analysis of SILAC methyl-proteomics data

To determine methylation sites that were significantly regulated in the SILAC experiments, MaxQuant results were processed as previously described in (Fong et al, 2019; Musiani et al, 2019; Musiani et al, 2020; Spadotto et al, 2020). Briefly, for each experiment, the mean and standard deviation of the unmodified peptides SILAC log2 ratios distribution were calculated; then, mean and standard deviation values were used to calculate the z-scores of the methyl-peptides; finally, a peptide was classified as significantly regulated in a given experiment if its z-score was > 2 or < −2 in both the FWD and the REV replicates of the experiment.

### Functional and structural analysis of the ProMetheusDB

Motif analysis was done using pLogo (O’Shea et al, 2013). Sequence windows centred on regulated (or unchanging) R-methyl-sites were submitted as foreground; sequence windows centred on all identified R-methyl-sites (including non-quantified ones) were submitted as background; the resulting position weight matrices were then downloaded and visualized with the Logomaker Python package (Tareen & Kinney, 2020). Functional enrichment analysis was performed using the “gprofiler2” R package (Kolberg et al, 2020); Gene Ontology terms, Reactome and KEGG pathways, and CORUM complexes were used as data sources; only the most significant, non-redundant terms are reported in the figures (full lists of enriched terms are reported in Table EV2). Overlap of the annotated R-methyl-sites with InterPro domains (Blum et al, 2021) was performed with an in-house Python script; the InterPro database was filtered to include only regions classified as “Domain” or “Homologous superfamily”. Mapping of modification sites on interaction surfaces was performed with the Mechismo web application (Betts et al, 2015), with a stringency threshold set to “medium”. The database of currently annotated phosphorylation sites was downloaded on 05-28-19 from Phosphosite Plus (Hornbeck et al, 2015). Protein:protein functional interaction networks were generated within Cytoscape (Shannon et al, 2003) using the Reactome plugin (Croft et al, 2011); protein:protein physical interactions were downloaded from the IMEX database (Orchard et al, 2012); network analysis was performed with Cytoscape and the Python package Pyntacle (Parca et al, 2020). Fisher’s exact tests were performed with the Scipy package in Python. Bar plots and box plots were generated with the Seaborn package in Python, network displays were generated in Cytoscape and the UpSets plot representation of methylation sites on proteins was generated with R.

### Protein immunoprecipitation (IP) of NONO and WB analysis of its R methylation state and co-IP of PSPC1

IP of NONO was performed starting from 1mg of HeLa whole-cell extract (WCE). Briefly, 30e6 HeLa cells were harvested, washed twice with cold PBS and re-suspended in 2 volumes of RIPA Buffer (10 mM Tris pH 8, 150 nM NaCl, 0.1 % SDS, 1 % triton, 1mM EDTA, 0.1% Na-Deoxycholate, 1mM PMSF, 1mM DTT and 1x Protease and Phosphatase Inhibitors cocktail (Roche), supplemented with 10 kU of Benzonase (Merck Life Science)). The suspension was rotated on a wheel for 45 min at RT (vortex every 10 min), centrifuged at 12.000 g for 1h at 4°C and the supernatant was transferred into a new Eppendorf tube. Proteins were quantified by BCA colorimetric assay (Pierce BCA Protein assay kit) and 1 mg of WCE was used for the IP, 8% of which was saved as Input (80 μg to be divided into 4 SDS-PAGE gels). The WCE used for the IP was rotated at 4°C overnight with 4μg of anti-NONO/p54 (sc-376865 Santa Cruz). G-protein-coupled magnetic beads (Dynabeads, Thermo Fisher Scientific) were saturated with a blocking solution (0.5% BSA) and rotated at 4°C overnight on a wheel. The following day, the beads were added to the lysate in 1: 100 proportion with the primary antibodies and incubated for 3 hours at 4°C on the wheel; the captured complexes were washed 4 times with the RIPA Buffer and then incubated 10 min at 95° with LSD sample Buffer (2X) supplemented with 100 mM DTT to elute the immunoprecipitated proteins. Equal protein amounts were separated by SDS-PAGE electrophoresis (Nupage® Novex® 4-12% Bis-tris Gel 1.5 mm, Thermo Fisher) and transferred on Transfer membrane (Immobilon-P, Merck Millipore) by wet-transfer method. Membrane blocking was performed with 10% BSA/TBS 0.1% Tween-20 for 1h at RT and followed by overnight incubation with the selected primary antibodies and subsequent incubation with the HRP-conjugated secondary antibodies (Cell Signaling Technology) for 1h at RT. Proteins were detected by ECL (Bio-Rad). The following primary antibodies were used:

- anti-NONO (SC-376865, 1:500) was purchased from Santa Cruz;
- anti-ADMA (ASYM24 07-414, 1:1000) and anti-SDMA (SYM10 07-412, 1:2000) were purchased from Millipore;
- anti-MMA (D5A12; 1:1000) was purchased from Cell Signaling Technology
- anti-PSPC1 (A302-461, 1:5000) were purchased from Bethyl Laboratories
- anti-GAPDH (Ab9484, 1:3000) was purchased from Abcam.

Quantification of the signal intensity for each band was performed by FiJi software (Schindelin et al, 2012) and signal intensity was normalized at 4 different levels:

- quantification of NONO in the Input was normalized on GAPDH (as loading control).
- quantification of NONO in its IP was normalized on the previous normalized input for each condition.
- R methylation (MMA, ADMA or SDMA) and PSPC1 signals were normalized on the amount of normalized NONO in the IP.
- signal intensity in MS023 condition was normalized on DMSO (untreated).

## Data Availability

The mass spectrometry proteomics data have been deposited to the ProteomeXchange Consortium via the PRIDE (Perez-Riverol et al, 2019) partner repository with the dataset identifier PXD027949.

## Acknowledgements

We thank all members of the Bonaldi group for their support, suggestions and critical discussion; in particular the ex-members of the group D. Musiani and V. Spadotto for producing part of the methyl-proteomics data that have been used to generate the ProMetheusDB. EM and MM are PhD students within the European School of Molecular Medicine (SEMM). LN was a PhD student within the SEMM.

## Authors contributions

EM developed hmSEEKER and drafted the manuscript; EM, FR and AY carried out the data analysis; MM and LN carried out the proteomics experiments; EM and AY trained the Machine Learning model; TB proposed the project, designed the experiments, defined the manuscript draft; TB and RG carried out the draft revision and evaluated the results; TB financially supported the project.

